# Inhibiting endothelial cell Mst1 attenuates acute lung injury in mice

**DOI:** 10.1101/2023.09.27.559864

**Authors:** Zhi-Fu Guo, Nopprarat Tongmuang, Chao Li, Chen Zhang, Louis Hu, Daniel Capreri, Mei-Xing Zuo, Ross Summer, Jianxin Sun

## Abstract

**Background:** Lung endothelium plays a pivotal role in the orchestration of inflammatory and injury responses to acute pulmonary insults. Mammalian sterile 20-like kinase 1 (Mst1), a mammalian homolog of Hippo, is a serine/threonine kinase that is ubiquitously expressed in many tissues and has been shown to play an important role in the regulation of apoptosis, inflammation, stress responses, and organ growth. While Mst1 exhibits high expression in the lung, its involvement in the endothelial response to pulmonary insults remains largely unexplored.

**Methods:** Mst1 activity was assessed in lung endothelium by western blot. Mst1 endothelial specific knockout mice and a pharmacological inhibitor were employed to assess the effects of Mst1 on homeostatic and lipopolysaccharide (LPS)-induced endothelial responses. Readouts for these studies included various assays, including NF-κB activation and levels of various inflammatory cytokines and adhesion molecules. The role of Mst1 in lung injury was evaluated in a LPS-induced murine model of acute lung injury (ALI).

**Results:** Mst1 phosphorylation was significantly increased in lung endothelial cells after exposure to tumor necrosis factor (TNF)-alpha (TNF-α) and mouse lung tissues after LPS exposure. Overexpression of full length Mst1 or its kinase domain promoted nuclear factor kappaB (NF-κB) activation through promoting JNK and p38 activation, whereas dominant negative forms of Mst1 (DN-Mst1) attenuated endothelial responses to TNF-α and interleukin-1β. Consistent with this, targeted deletion of Mst1 in lung endothelium reduced lung injury to LPS in mice. Similarly, wild-type mice were protected from LPS-induced lung injury following treatment with a pharmacological inhibitor of Mst1/2.

**Conclusions:** Our findings identified Mst1 kinase as a key regulator in the control of lung EC activation and suggest that therapeutic strategies aimed at inhibiting Mst1 activation might be effective in the prevention and treatment of lung injury to inflammatory insults.

## Introduction

Acute lung injury (ALI) and its more severe form, the acute respiratory distress syndrome (ARDS) are inflammatory lung disorders that cause significant morbidity and mortality in a considerable proportion of hospitalized patients ^1^. The lung endothelium has been identified as an essential player in orchestrating and propagating ARDS ^2^, and disruption of lung endothelium is one of the major hallmarks of ARDS pathobiology ^3 4^. Under normal physiological conditions, the lung endothelium exhibits an inhibitory effect on inflammation ^5 6^. However, in response to a wide range of pathological insults, such as hypoxia, bacterial toxins, and stimulation of cytokines and chemokines, activation of the lung endothelium leads to the recruitment of inflammatory cells to vascular walls and the transmigration of leukocytes into the lung ^7^. As such, therapeutic strategies aimed at reducing endothelial activation are now believed to be important for decreasing the onset and severity of ARDS and other inflammatory vascular conditions ^8 9^.

Activation of the transcription factor nuclear factor (NF)-κB plays an important role in mediating expression of adhesion molecules and the subsequent interaction of immune cells with endothelial cells ^10 11^. The promoter regions of genes for adhesion molecules and cytokines normally have binding sites for NF-κB. In resting ECs, NF-κB resides as an inactive molecule in the cytoplasm by forming complexes with inhibitors, such as the IκB proteins (IκBs). However, upon stimulation by proinflammatory cytokines, IκBs are phosphorylated by IκB kinase (IKK), ubiquitinated, and proteolytically degraded by 26S proteasomes, which allows NF-κB to translocate to the nucleus, where it can bind to promoter regions and induce the transcription of key inflammatory target genes, such as a host of adhesion molecules, inflammatory cytokines, and chemokines ^11^. Notably, several therapeutic approaches to suppress inflammation are based on the ability to control NF-κB activation, although these have yet to prove effective in patients ^12 4^. Thus, further understanding the mechanisms underlying NF-κB activation will help for developing novel therapies for inflammatory vascular diseases, such as acute lung injury and ARDS.

Mammalian sterile 20-like kinase 1 (Mst1), a mammalian homolog of Hippo, is a ubiquitously expressed serine/threonine kinase that has been implicated in regulating various cellular processes, including cell proliferation, apoptosis, and immune responses^13 14^. Indeed, in response to various stimuli and cellular stresses, including TNF-α, serum starvation, staurosporine, UV irradiation, as well as anti-cancer drugs, Mst1 is activated in a number of cell lines, including BJAB, 293T, COS-1, Jurkat, HeLa and HL-60 ^14^. Furthermore, Mst1 has been shown to be activated in response to ischemia/reperfusion in cardiac myocytes and other types of vascular injury ^15 16 17 18^, further suggesting its important roles in the pathogenesis of cardiovascular diseases. Inflammation, apoptosis, and cellular stress play key roles in tissue damage of ALI and ARDS. Although Mst1 is highly expressed in the lung ^19 20^, its involvement in regulating signaling pathways of lung injury remains largely unknown. Investigating how Mst1 influences inflammatory responses in the lungs could provide valuable insights into the underlying mechanisms and potential therapeutic strategies for this condition.

In this study, we hypothesized that Mst1 might be involved in regulating endothelial inflammatory responses to pulmonary insults. To test this hypothesis, we employed both genetic and pharmacological approaches to systematically dissect the role of Mst1 in lung endothelial biology. In brief, our findings indicate that Mst1 is a critical regulator of inflammatory response in lung endothelium. More specifically, we show that Mst1 activation promotes endothelial inflammation and worsens lung injury to LPS whereas targeted deletion of Mst1 in lung endothelium or pharmacological inhibitors of Mst1/2 effectively mitigate LPS-induced inflammatory responses in the lung.

## Methods

### Cell Culture

Human lung microvascular ECs were purchased from ATCC and cultured in EBM-2 medium supplemented with EGM-2 BulletKit (Lonza). THP-1 and U937 cells were purchased from ATCC and cultured in RPMI medium 1640 (Corning, 10-040-CM). EA.hy926 cells were purchased from ATCC and cultured in Dulbecco Modified Eagle Medium (DMEM, Corning, 10013CV) DMEM). All medium was supplemented with fetal bovine serum (FBS) (Gibco, 1082147) and penicillin/streptomycin (Corning, 30-002-CI).

### Antibodies and Reagents

Anti-Mst1 (#14946, 1:500 dilution), anti-p65 (#8242, 1:1000 dilution), anti-p38 (#9212, 1:1000 dilution), anti-pjospho-p38 (#9211, 1:1000 dilution), anti-Erk1/2 (#4695, 1:1000 dilution), anti-phospho-Erk1/2 (#4370, 1:1000 dilution), anti-JNK (#9252, 1:1000 dilution), anti-phospho-JNK (#9251, 1:1000 dilution), antianti-p-MOB1 (#8699, 1:1000 dilution), and anti-MOB1 (#13730, 1:1000 dilution) antibodies were purchased from Cell Signaling Technology. Anti-phospho-Mst1 (AP0906, 1:500 dilution) antibody was acquired from ABclonal Technology. Anti-ICAM1 (sc-7891, 1:800 dilution), anti-VCAM1 (sc-13160, 1:300 dilution), and anti-GAPDH (sc-32233, 1:1000 dilution) antibodies were from Santa Cruz Biotechnology. IRDye 680RD Donkey anti-Rabbit (926-68073, 1:10,000 dilution) and 800CW Goat anti-Mouse (926-32210, 1:10,000 dilution) antibodies were obtained from LI-COR Bioscience. LPS (sc-3535) was acquired from Santa Cruz Biotechnology. Recombinant Human TNF-α (300-01A) and IL-1β (200-01B) were acquired from PeproTech. DAPI (4’,6-diamidino-2-phenylindole) (D1306) was purchased from Invitrogen. XMU-MP1 (#22083, Cayman) was dissolved in 2% DMSO+30% PEG 300+2% Tween 80+ddH2O at 5mg/mL for in vivo studies. JNK inhibitor (SP600125, #HY-12041) and p38 MAPK inhibitor (SB203580, #HY-10256) were purchased from MedChemExpress.

### Mice

Mst_1/2_ floxed mice (STK3^f/f^/STK4^f/f^ mice were purchased from The Jackson Laboratory (Stock # 017635, JAX) and crossed with C57BL/6J to generate Mst1 floxed mice (Mst1^f/f^) mice with exons 4 and 5 of Mst1 flanked by loxP sites^21^. Mst1^f/f^ mice were then interbred with Tie2-Cre mice (Stock number: 008863, the Jackson Laboratories) to generate endothelial specific Mst1 knockout mice (Mst1^ΔEC^). All mice were genotyped by PCR. The genomic DNA isolated from mice tail biopsies was analyzed by PCR to detect Mst1 WT allele (product size 303 bp), loxP-flanked allele (450 bp), and Tie2-Cre allele (100bp). Genotyping primers for Mst1 allele were 5’-AGT GTT GGC TCT TGA TTT TCC T-3’ (forward) and 5’-CAG GGC TAG AGT GAA ACC TTG-3’ (reverse), and genotyping primers for Cre allele were 5’-GCGGTCTGGCAGTAAAAACTATC-3 ′ (forward) and 5’-GTGAAACAGCATTGCTGTCACTT-3’ (reverse). All mice were on the C57BL/6J background and maintained under specific pathogen–free conditions at 22°C with a 12-hour light/12-hour dark cycle. Age-matched littermates (8–12 weeks old) were used for the experiments. All animal protocols were approved by the Institutional Animal Care and Use Committee at Thomas Jefferson University before performing any studies.

#### Murine ALI model

ALI was induced as described previously ^22^. In brief, anesthetized mice (8–12 wk, 20–25 g) were suspended from a sloped board, and a one-time dose of LPS (100 μg per mouse) was instilled in posterior oropharyngeal space. For preventive studies, XMU-MP1 (4mg/kg, IP) was administrated 2 hours before LPS instillation. For treatment studies, XMU-MP1 (10mg/kg, IP) was administrated at 0.5 h and 12 h after LPS instillation. 24 h after LPS administration, lung tissues were harvested for further analysis. Researchers were blinded to the assignment of treatment and evaluation of the experimental results.

#### Bronchoalveolar lavage

Bronchoalveolar lavage (BAL) was performed by cannulating the trachea with a blunt 22-gauge needle and instilling the same 1 ml of sterile PBS back and forth in the airways three times as described previously ^22^. Total cell counts were performed using a TC20 automated cell counter (Bio-Rad Laboratories, Hercules, CA), and differential cell counts were performed after cytocentrifuging cells on glass slides and staining with HEMA 3 (Fisher Scientific, Tustin, CA). Total protein levels were determined as previously described ^22^. The activity of the enzyme myeloperoxidase (MPO), a marker of polymorphonuclear neutrophil primary granules, was determined in a lung fragment (upper right lobe) after pulmonary lavage as described previously ^23^.

#### Enzyme-linked immunosorbent assay

MCP-1, TNF-α, and IL-6 levels were quantified in BALF and lung tissues using commercially available DuoSet ELISA kits (R&D Systems) according to manufacturer’s instructions and published protocols ^22^. Briefly, Nalgene Nunc Maxisorp plates were coated overnight with antibodies to MCP-1 (4 μg/ml), TNF-α (4 μg/ml), and IL-6 (4 μg/ml) and the next morning plates were washed and blocked for 2 hours. Samples were added to the wells at various dilutions, followed by incubation with detection antibody for 2 hours. Plates were subsequently washed, and streptavidin-horseradish peroxidase conjugate antibody was added to each well for 20 min. This was followed by an additional wash step, before plates were incubated with 3,3′,5,5′-tetramethylbenzidine (TMB) substrate solution (R&D Systems). Enzymatic reactions were quantified by measuring absorbance at 450 nm using a standard plate reader (Biotek Instrument).

#### Lung histology

Lung histology was performed on paraformaldehyde-fixed tissues embedded in paraffin wax. Sections (5 μm) were placed on positively charged glass slides, and tissues were deparaffinized before undergoing hematoxylin and eosin (H&E) staining. Tissues were visualized using standard light microscopy ^24^.

#### Pulmonary microvascular permeability

Lung permeability was assessed by quantifying the extravasation of Evans blue dye (E2129, Sigma-Aldrich). Evans blue (20 mg/kg; Sigma-Aldrich) was injected intravenously 2 hours before euthanasia. Before lungs were extracted from the thorax, the pulmonary vasculature was gently perfused by slowly injecting saline in the spontaneously beating right ventricle. Extraction of Evans blue dye was performed by incubating tissues at 65°C with formamide (2 ml/g tissue) overnight. Lung tissues were then centrifuged (12,000 g for 30 min), and Evans blue dye concentration in supernatant was determined spectrophotometrically by measuring absorption at 620 nm and correcting for the presence of heme pigments: A620 (corrected) = A620 − (1.426 × A740 − 0.030) where A620 is absorbance at 620 nm and A740 is absorbance at 740 nm.

### Murine Lung EC Isolation

The lungs from WT and Mst1-KO mice (at 7–9 d of age, three per group) were dissected into single lobes and incubated in Dulbecco’s modified Eagle medium containing collagenase solution. The cell suspension was purified with anti-CD31–coated magnetic beads (Invitrogen) and cultured in extracellular matrix medium (ScienCell) supplemented with 1% penicillin–streptomycin solution, 1% EC growth stimulant, and a 20% FBS BulletKit (Lonza) as described previously ^24^.

#### Monocyte adhesion assay

Lung ECs transfected with Ad-LacZ or Ad-DNMst1 were treated with 10 ng/ml IL-1β or 20 ng/ml TNF-α for 8 hours to induce the expression of adhesion molecules before adhesion assay. THP-1 cells were labeled with calcein-AM (Invitrogen) according to the manufacturer’s instruction. After lung ECs were stimulated and washed, 2.5×10^5^ calcein-labeled THP-1 cells were added to each well and allowed to interact for 60 min at 37°C as we described previously^25^. Unbound cells were removed by gently washing with complete medium, and the number of attached THP-1 cells was counted on an inverted fluorescent microscope.

#### Immunofluorescence staining

Lung ECs were fixed and sequentially incubated with primary antibodies and appropriate fluorescent-labeled secondary antibodies. Images were visualized using an Olympus IX70 epifluorescence microscope as previously described ^26^.

#### Construction of adenoviruses

Adenoviruses harboring wild-type Mst1 (Ad-Mst1) and dominant-negative Mst1 (Ad-DN-Mst1) were made using AdMax (Microbix) as previously described ^27^. The viruses were propagated in AD-293 cells and purified using CsCl2 banding, followed by dialysis against 20 mmol/L Tris-buffered saline with 10% glycerol. Titering was performed on Ad293 cells using Adeno-X Rapid Titer kit (Clontech) according to the instructions of the manufacturer ^28^.

#### Quantitative Real Time PCR (qRT-PCR)

Total RNA was extracted from human lung ECs and mouse lung tissues using TRIZOL reagent kit (Invitrogen). The cDNA was generated using iScript cDNA synthesis kit (170-8891; Bio-Rad). Real-time PCR analysis was performed with Power SYBR Green PCR Master Mix (4367659; Life Technologies) by a StepOne Plus system (Applied Biosystems). GAPDH RNA was used as an internal control. The relative difference was expressed as the fold matched control values calculated with the efficiency-corrected 2^−ΔΔCt^ method. The primer list for quantitative RT-PCR is provided in the Supplemental Table I.

#### Western Blot Analysis

Western blot analysis was performed as we described previously ^26^. In brief, cell or tissue lysates were resolved by SDS-PAGE and transferred to nitrocellulose membrane (Bio-Rad Laboratories, USA). Blots were blocked with 5% nonfat milk in phosphate-buffered saline (PBS), then developed with 5% PBST (PBS with 0.1% Tween20) and diluted first antibodies followed by incubating with either IRDye 700 or 800 labelled secondary antibodies and then visualized using Odyssey Infrared Imaging System (Li-Cor, Lincoln, NE). The intensity of the bands was quantified by using the Odyssey software.

#### Luciferase Reporter Assay

Endothelial cell line EA.Hy926 cells were seeded in 12-well plates and incubated overnight. The cells were transfected with 100 ng of NF-κB firefly luciferase reporter plasmid p(NF-κB)_3_-Luc (Stratagene) and 10 ng of Renilla luciferase reporter plasmid pRL-RSV (Promega), in the presence or absence of indicated expression vectors using FuGene 6 transfection reagent (Promega). 36 hours after transfection, cells were treated with or without indicated stimuli for 8 hours, then directly lysed in the lysis buffer (Promega). Cell lysates (20 μL) were assayed for luciferase activity using the dual Luciferase Assay System (Promega, USA), according to manufacture instructions. We used Renilla luciferase as a control for transfection efficiency. Firefly luciferase activity was normalized for transfection efficiency by corresponding Renilla luciferase activity ^25^.

#### Electrophoretic Mobility Shift Assay (EMSA)

EMSA was performed as previously described ^29^. Briefly, the oligonucleotides corresponding to the consensus sequence of NF-κB (5′-AGTTGAG<u>GGGACTTTCCC</u>AGGC-3′), AP1 (5′-CGCTTGA<u>TGACTCA</u>GCCGGAA-3′) and Nur77 (5′-CTAGCGCT<u>TGACCTTT</u>TCGCACGAAGATC-3′) were synthesized and labeled with IRDye 700 (IDT). EMSA were performed with Odyssey® IRDye® 700 infrared dye labeled double-stranded oligonucleotides coupled with the EMSA buffer kit (Thermo Scientific) as we described previously ^26,29^.Briefly, 5 µg of nuclear extract was incubated with 1 µl of IRDye® 700 infrared dye labelled double-stranded oligonucleotides, 2 µl of 10 × binding buffer, 2.5 mM DTT, 0.25% Tween-20 and 1 µg of poly (dI-dC) in a total volume of 20 µl for 20 min at room temperature in the dark. The specificity of the binding was examined using competition experiments, where 100-fold excess of the unlabeled oligonucleotides were added to the reaction mixture for 30 minutes. The gel supershift assay was performed by adding p-65 antibody (Cell Signaling) prior to the addition of the fluorescently labeled probe. Sample proteins were separated on a 4% polyacrylamide gel in 0.5 × Tris-borate-EDTA running buffer for 60 min at 70V. The gel was scanned by direct infrared fluorescence detection on the Odyssey® imaging system (LI-COR Bioscience).

#### Statistical Analysis

Data were presented as the mean ± SD. Normality of distribution was assessed by using Shapiro-Wilk test. Differences between groups with normally distributed data were analyzed using a two-sided unpaired t test (for 2 groups of data), one-way ANOVA followed by Tukey post hoc test (for 3 or more groups of data). Two-way ANOVA coupled with Tukey’s post hoc test was applied for two independent variables. For non-normally distribution or sample number less than 6, two-tailed Mann-Whitney test (2 groups) or Kruskal-Wallis test (multiple groups) was performed, followed by Dunn post hoc analysis. Values of *P*<0.05 were considered statistically significant.

## Results

### Mst1 is activated in lung ECs after exposure to TNF-α

To investigate the role of Mst1 in pulmonary vascular inflammation, we examined the activation and expression of Mst1 in lung ECs in response to TNF-α, which is a recognized inducer of endothelial activation and plays an important role in the pathobiology of ALI and ARDS in humans^30 31^. The activation of Mst1, as indicated by its phosphorylation at Thr183 ^32^, was determined by western blot at different time points after TNF-α stimulation. As shown in Figure 1A, TNF-α stimulation significantly increased Mst1 phosphorylation at Thr183 in a time dependent manner. The maximal induction of TNF-α on Mst1 phosphorylation was observed at 20 mins, when Mst1 phosphorylation was increased by approximately 2.5-fold. Mst1 is highly expressed in endothelial cells in lung tissues (Figure S1). To determine whether Mst1 activation is involved in inflammatory lung diseases, we generated a mouse model of ALI induced by LPS instillation ^24^. As shown in Figure 1B, Mst1 phosphorylation, as determined by western blot, was significantly increased in a time dependent manner in mouse lung tissues after LPS instillation. Together, these data suggest that Mst1 may regulate inflammatory responses in lung ECs.

**Figure 1.**
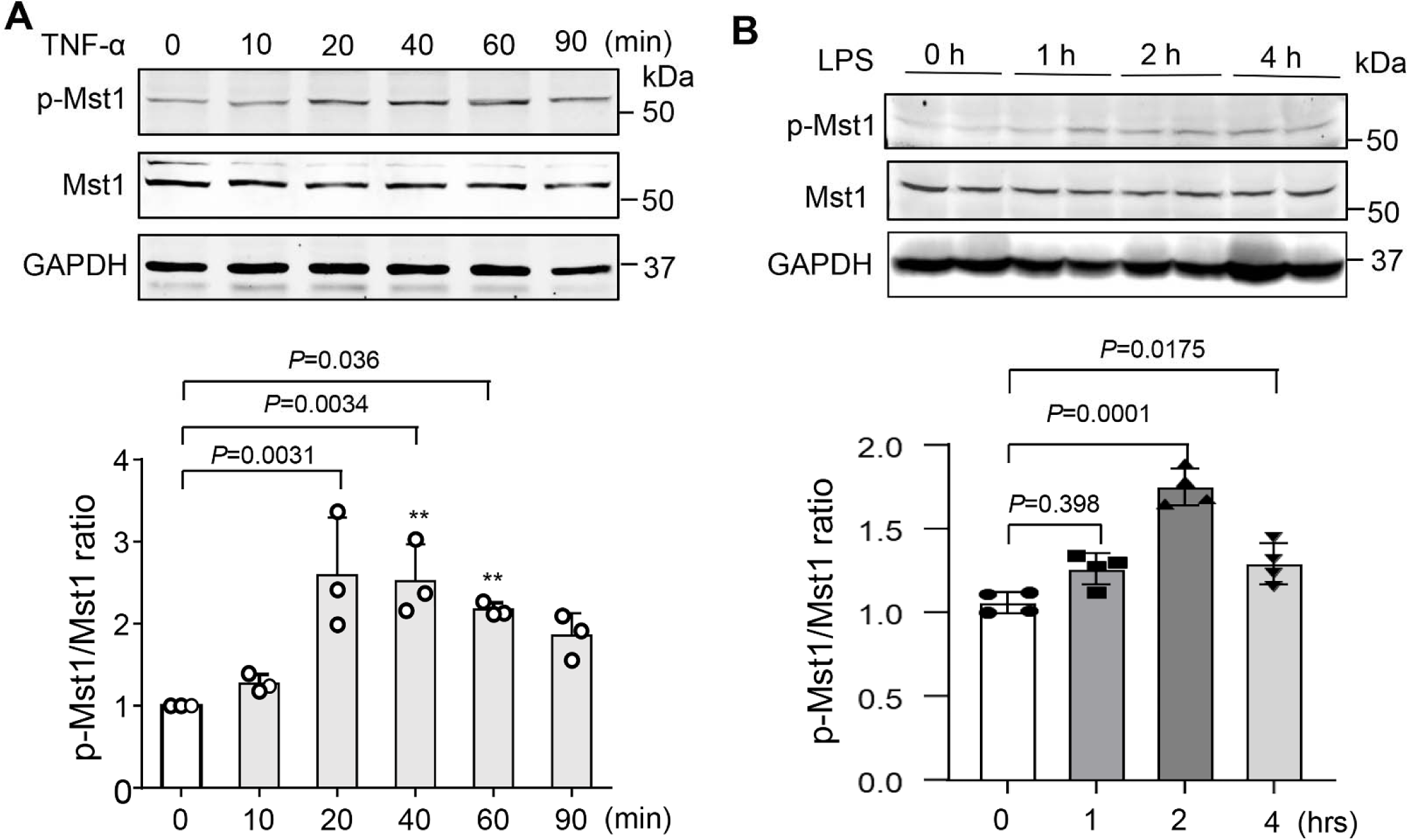
TNF-α induces Mst1 activation in lung ECs. **A**, Lung ECs were treated with 20 ng/ml TNF-α at the indicated time points. Phosphorylation and total protein levels were determined by western blot and results were quantified by densitometry analysis. n = 3. Statistical significance was determined by 1-way ANOVA with Sidak multiple comparisons. **B**, Mice were subjected to LPS instillation. Lung tissues were collected for the determination of Mst1 phosphorylation by western blot. n = 4. Statistical significance was determined by a 1-way ANOVA with Sidak multiple comparisons (**A** and **B**)

### Mst1 induces NF-κB Activation

Activation of the NF-κB signaling is essential for driving many inflammatory and immunological responses^10^. To determine whether Mst1 plays a role in the regulation of pulmonary vascular inflammation, we examined the effect of Mst1 overexpression on NF-κB promoter activity at the transcriptional level in ECs. Mst1 contains an N-terminal kinase domain (aa 1–325), inhibitory domain (aa 326–294), and a C-terminal dimerization domain (aa 395–487) ^28,33^ (Figure 2A). We generated a series of Mst1 deletion mutants and transfected these mutants into the endothelial cell line Hy.EA296 cells along with a reporter plasmid containing a heterologous promoter driven by NF-κB elements upstream of the luciferase gene. As shown in Figure 2B, transfection of the full length and Mst1 mutants containing its kinase domain significantly augmented the NF-κB promoter activity, whereas transfection of Mst1 dominant negative mutant (K59R, DN-MST1) ^28^, C-terminal inhibitory domain, and dimerization domain had little to no effect on the basal NF-κB promoter activity. Importantly, overexpression of DN-Mst1, which has been shown to inhibit endogenous Mst1 activity in various types of cells ^15 28^, markedly suppressed TNF-α–induced NF-κB activation, as measure by both luciferase activity and EMSA (Figure 2C and 2D). MST1 is an upstream kinase of the JNK and p38 MAPK ^34^, which are involved in the activation of NF-κB. In human lung ECs, we found that Mst1 overexpression induced phosphorylation of JNK and p38 MAPK (Figure 2E), and pharmacological inhibition of JNK and p38 MAPK significantly attenuated Mst1-induced NF-κB activation in ECs (Figure 2F). Together, these results suggest that Mst1 induces NF-κB activation at least in part through targeting the JNK and p38 MAPK in lung ECs.

**Figure 2.**
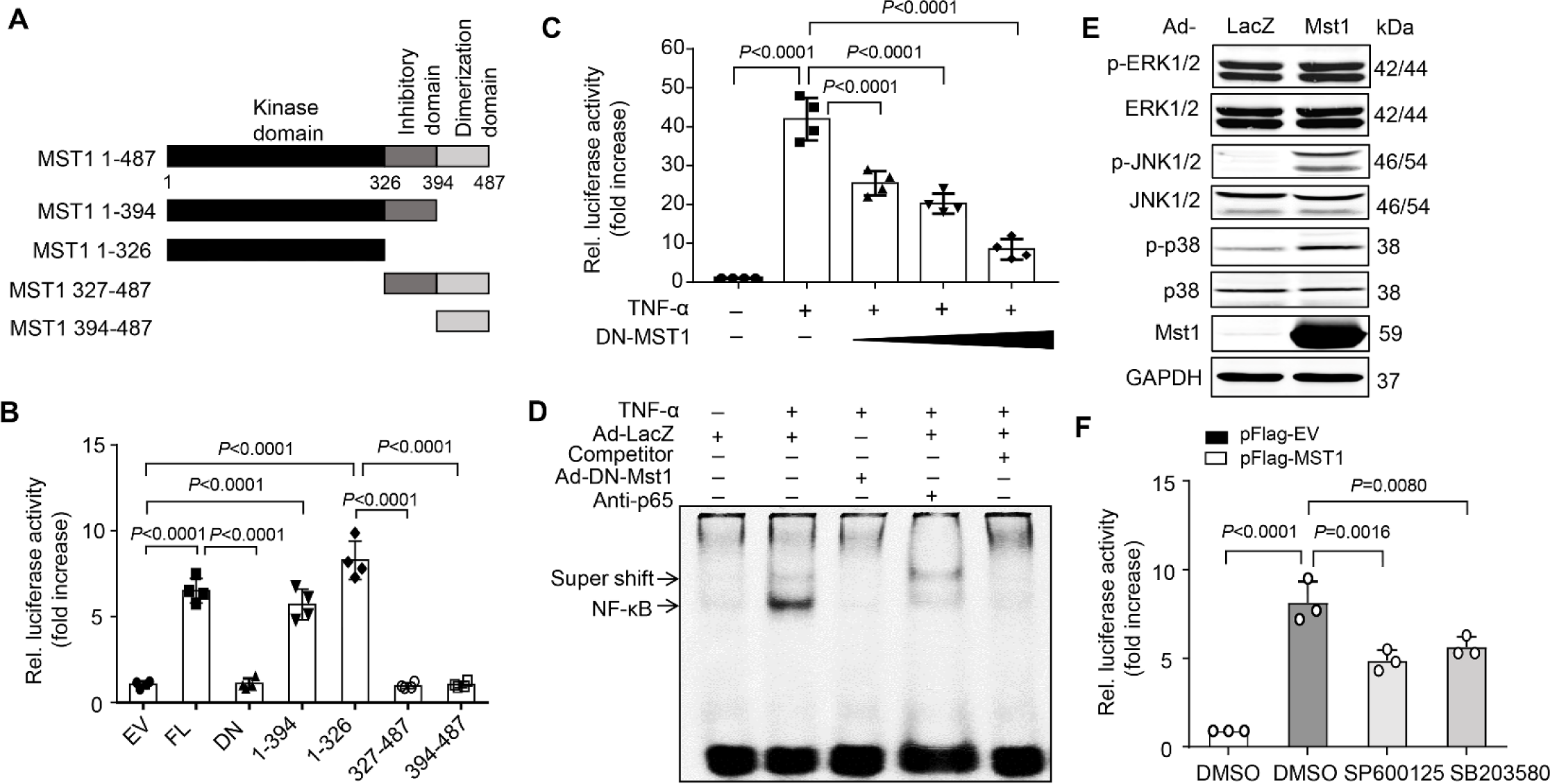
Mst1 induces NF-κB activation in ECs. **A**, Schematic representation of Mst1 mutants. **B**, EA-Hy926 cells were transfected with 100 ng of p(NF-κB)3-Luc, 10 ng of pRL-RSV, and 200 ng of either empty vector (EV) or Mst1 mutants. Thirty-six hours after transfection, luciferase assays were performed (n=4). **C**, Lung ECs were transfected with 100 ng of p(NF-κB)3-Luc, 10 ng of pRL-RSV, and increasing amounts of pFlag-DN-Mst1. Thirty-six hours after transfection, luciferase assays were performed 6 hours after treatment with or without 20 ng/mL TNF-α (n=4). **D**, Lung ECs were transduced with adenoviruses bearing LacZ (Ad-lacZ) or DN-Mst1 (Ad-DN-Mst1) (mois, 50) for 48 hours and then treated in the presence or absence of 20 ng/mL TNF-α for 1 hour. Nuclear protein was extracted, and EMSA was performed. The NF-κB complex was partially supershifted by anti– NF-κB p65 antibody or blocked by cold competitive probe. **E**, Human lung ECs were transduced with Ad-LacZ or Ad-Mst1 (mois, 50). 48 hours after transduction, cell lysates were collected for western blot analysis. **F**, EA-Hy926 cells were pretreated with DMSO vehicle control, JNK inhibitor (SP600125, 20 µmol/L) or p38 MAPK inhibitor (SB203580, 20 µmol/L) for 1 h. Subsequently, the cells were transfected with 500 ng of p(NF-kB)3-Luc, 50 ng of pRL-RSV, 200 ng of either empty vector (pFlag-EV) or pFlag-Mst1. 24 hours after transfection, luciferase assays were performed using Dual-Luciferase® Reporter Assay System. Significance was determined by a 2-way ANOVA with Bonferroni’s post hoc test (**B**, **C**, and **F**).

### Inhibition of Mst1 Suppresses cytokine-Induced lung EC activation

Activation of NF-κB is responsible for EC activation and the cytokine-induced expression of adhesion molecules, such as VCAM-1 and ICAM-1, which mediate the adhesion of monocytes to inflamed ECs ^25^. Thus, we examined the effects of Mst1 inhibition on the TNF-α– and IL-1β–induced-induced expression of inflammatory molecules in human lung ECs. As shown in Figure 3A, adenovirus-mediated overexpression of DN-Mst1 substantially attenuated the TNF-α– and IL-1β–induced-expression of MCP-1, IL-6, ICAM-1, and VCAM-1, as determined by qPCR. Western blot demonstrated that both TNF-α– and IL-1β–induced-expression of ICAM-1 and VCAM-1 were also markedly suppressed in DN-Mst1 overexpressing ECs (Figure 3B). Monocyte adhesion to ECs is an important event in the initiation of vascular inflammation ^25^. Therefore, we examined the effect of overexpressing DN-Mst1 on THP-1 cell adhesion to the activated lung ECs. Stimulation of lung ECs with either TNF-α (20 ng/ml) or IL-1β (10 ng/mL) substantially increased THP-1 adhesion, which was markedly suppressed by approximately 85% in response to overexpressing DN-Mst1 (Figure 3C). Taken together, these results further suggest that Mst1 functions as a positive regulator of the cytokine-induced inflammatory responses in lung ECs.

**Figure 3.**
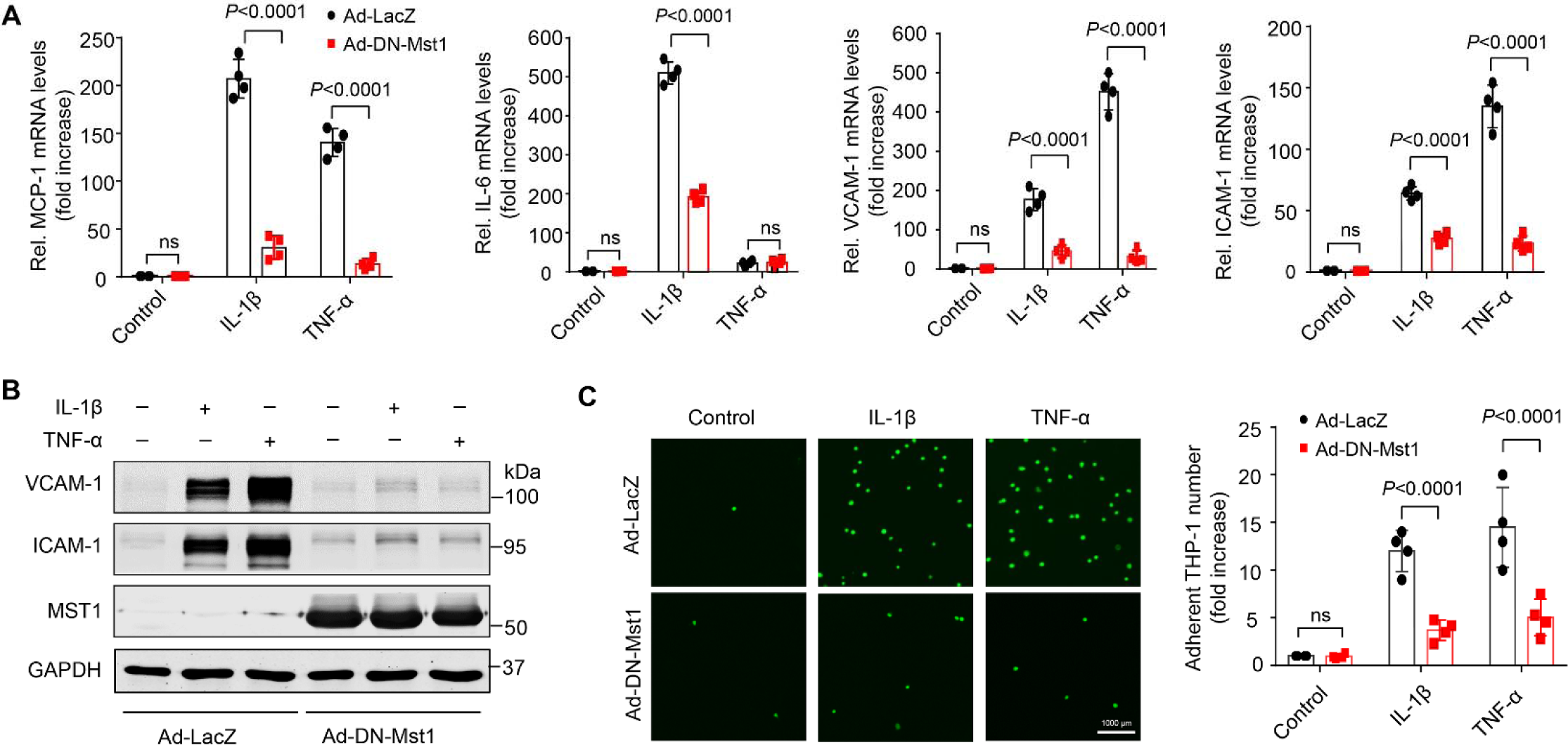
Overexpression of DN-Mst1 attenuates lung EC activation. Lung ECs were transduced with Ad-LacZ, Ad-Mst1, or Ad-DN-Mst1 (mois, 50). 48 hours after adenovirus transduction, ECs were treated with or without 20 ng/mL TNF-α or 10 ng/mL IL-1β for 8 hours. The expression of MCP-1, IL-6, VCAM-1, and ICAM-1 was detected using qPCR (**A**) and by western blot (**B**) using anti-VCAM-1 or anti-ICAM-1 antibodies. **C**, DN-Mst1 inhibits THP1 adherences to lung ECs. ECs were transduced with Ad-LacZ or Ad-Mst1 (mois, 50). Forty-eight hours after adenovirus transduction, lung ECs were treated in the presence or absence of 20 ng/mL TNF-α or 10 ng/mL IL-1β for 8 hours and incubated with calcein-labeled THP1 cells for another 1 hour. Following washing, attached THP1 cells were visualized and counted with an inverted fluorescent microscopy. Scale bars 1000 μm. n=4. Significance was determined by a 2-way ANOVA with Bonferroni’s post hoc test (**A** and **C**). ns indicates nonsignificant.

### Mst1 deletion in ECs attenuates LPS-induced acute lung injury in mice

To define the *in vivo* functional significance of Mst1 in EC activation, we generated EC–specific Mst1 knockout mice (Mst1^fl/fl^/Cre^+/–^, Mst1^ΔEC^) by crossing Mst1^fl/fl^ mice (Mst1^wt^) with Tie2-Cre transgenic mice (Figure 4A). We observed Mst1^ΔEC^ mice for one year and did not appreciate any detectable phenotypic differences with Mst1^wt^mice. To further verify the KO efficacy of Mst1 in vasculature, we determined Mst1 expression by qPCR in isolated lung ECs of Mst1^wt^ and Mst1^ΔEC^ mice. As shown in Figure 4B, Mst1 mRNA levels in lung ECs were significantly decreased by nearly 85% in Mst1^ΔEC^ mice as compared with Mst1^wt^ mice. To test the role of endothelial Mst1 in acute lung injury responses, we instilled LPS in the airways of Mst1^wt^ and Mst1^ΔEC^. To quantify lung injury responses in these mice, we first measured the total number of inflammatory cells and total protein levels in the BALF. As shown in Figure 4C, endothelial specific depletion of Mst1 markedly attenuated the total number of inflammatory cells and protein levels in the BALF after LPS exposure. Furthermore, as shown in Figure 4D and E, neutrophil infiltration and the expression of cytokines and adhesion molecules were substantially reduced in the lungs of Mst1^ΔEC^ compared with Mst1^wt^. Together, these data suggest that endothelial Mst1 plays a pivotal role in LPS-induced acute lung injury in mice.

**Figure 4.**
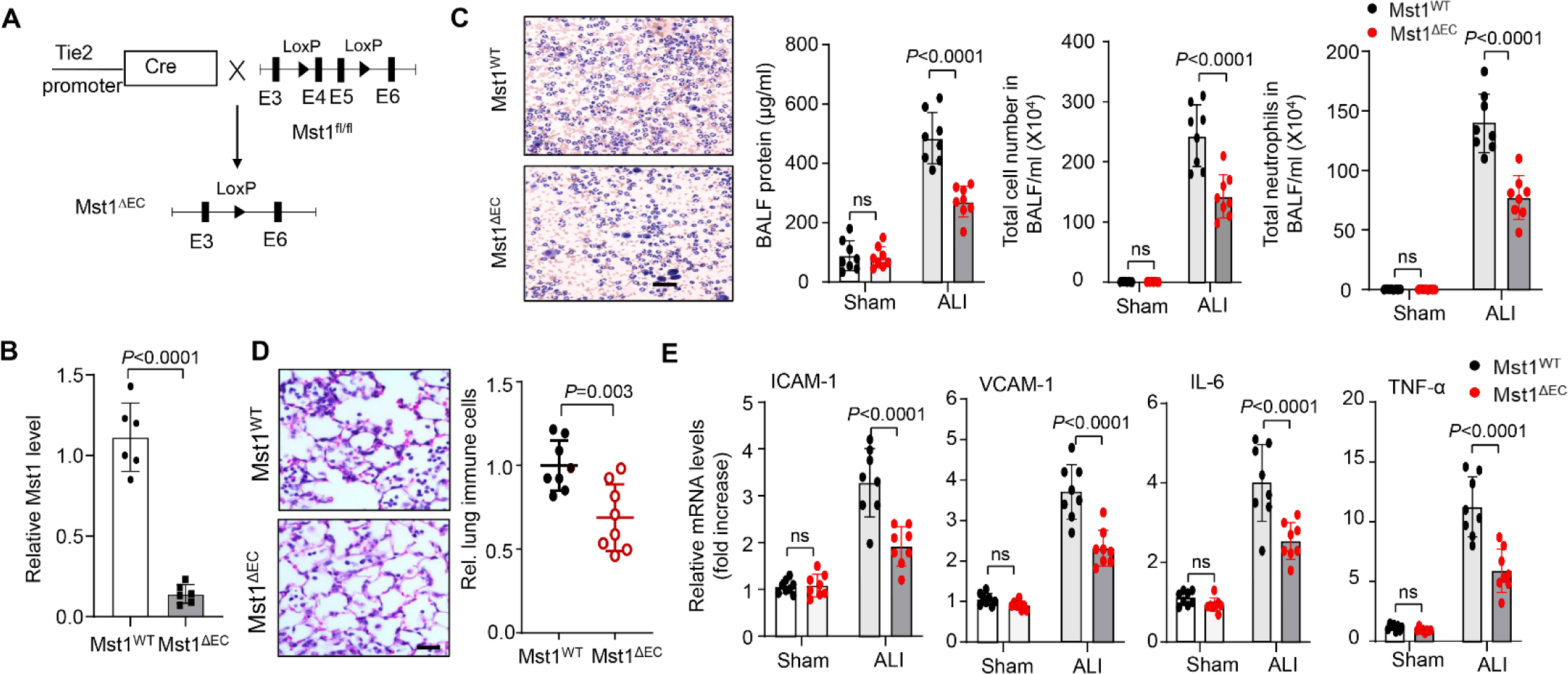
Endothelial specific deficiency of Mst1 attenuates LPS-induced lung injury in mice. **A**, Strategy of generating Mst1 endothelial KO mice. **B**, Expression of Mst1 in lung ECs isolated from Mst1^WT^ and Mst1^ΔEC^ mice was determined by RT-PCR. n = 6. **C**, Total cell and neutrophil counts and protein levels in bronchoalveolar lavage fluid (BALF) of Mst1^WT^ and Mst1^ΔEC^ mice after LPS instillation (*n* = 8). **D**, Mst1^WT^ and Mst1^ΔEC^ mice were subjected to LPS instillation. Lung samples was harvested from mice at 24 hours after treatment. Representative hematoxylin and eosin staining (n = 8 mice per group) of lung sections shows marked suppression of lung inflammatory injury in Mst1^ΔEC^ mice compared with Mst1^WT^ mice. Scale bars, 100 μm. n=8. **E**, Mst1^WT^ and Mst1^ΔEC^ mice were subjected to LPS instillation. Lung samples was harvested from mice at 24 hours after LPS treatment. Expression of cytokines and adhesion molecules was determined by qPCR. n=8. Significance was determined by a 2-way ANOVA with Bonferroni’s post hoc test (**C** and **E**) and a Student’s t test (**D**). ns indicates nonsignificant.

### Pharmacological inhibition of Mts1 attenuates lung EC activation and ALI in mice

To further explore the therapeutic capacity of targeting Mst1 for the treatment of ALI, we tested whether pharmacological inhibition of Mst1 could attenuate ALI in mice. Since specific inhibitors of Mst1 are not available, we examined the effects of XMU-MP1 (Figure 5A), which has been shown to selectively block both Mst1 and Mst2 ^35^, on endothelial activation. We first examined the inhibitory effects of XMU-MP1 on Mst1 in ECs, as determined by measuring phosphorylation of its substrate MOB1 (Mps One Binder 1) by western blot. As shown in Figure 5B, XMU-MP1 dose-dependently inhibited phosphorylation of MOB1 in human lung ECs with IC_50_ of 1.591 μmol/L. We then determined the effects of XMU-MP1 on expression of cytokines and adhesion molecules in ECs. To this end, we pretreated human lung ECs with 2 μmol/L XMU-MP1 for 2 hours before the stimulation of ECs with TNF-α for 12 hours. We then determined the expression of cytokines and adhesion molecules by qPCR. As shown in Figure 5C, TNF-α-induced increased expression of MCP-1, IL-6, and VCAM-1 was significantly inhibited by XMU-MP1, whereas ICAM-1 mRNA levels were not similarly affected. However, XMU-MP1 significantly inhibited TNF-α-stimulated protein expression of VCAM-1 and ICAM-1, with IC50 of 1.0828 μmol/L and 1.7637 μmol/L respectively, and this associated with marked suppression of monocyte adhesion to activated ECs and reduced transendothelial migration (Figure 5D and E and Figure S2).

**Figure 5.**
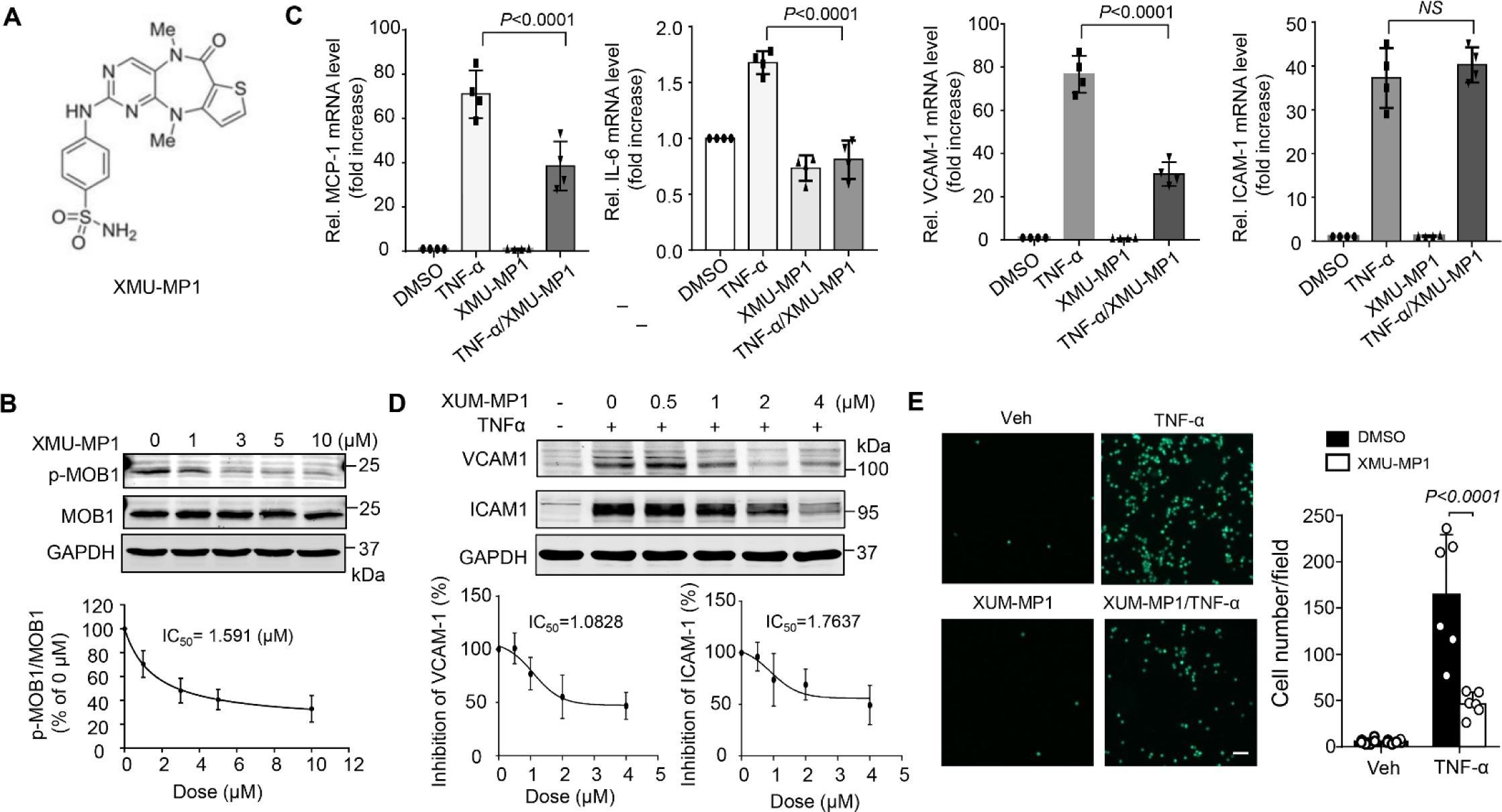
Pharmacological inhibition of Mst1 attenuates lung EC activation. **A**, The structure of Mst1/2 inhibitor XMU-MP1. **B**, Human lung ECs were treated with indicated concentrations of XMU-XP1 for 24 hours. Phosphorylation of MOB1 was determined by western blot. **C**, Lung ECs were pretreated with vehicle (DMSO) or 2 μmol/mL XMU-MP1 for 2 hours and then stimulated with TNF-α for 6 hours. The expression of MCP-1, IL-6, VCAM-1, and ICAM-1 was determined by qPCR. n=4. Significance was determined by two-way ANOVA with Bonferroni’s post hoc test. **D**, Lung ECs were pretreated with vehicle (DMSO) or increasing concentration of XMU-MP1 for 2 hours and then stimulated with TNF-α for 12 hours. The expression of VCAM-1 and ICAM-1 was determined by western blot. The expression of VCAM-1 was quantitated by densitometric analysis. IC_50_ was then calculated. n=3. **E**, Lung ECs were pretreated with vehicle (DMSO) or 2 μmol/mL XMU-MP1 for 2 hours and then treated in the presence or absence of 20 ng/mL TNF-α for 8 hours and incubated with calcein-labeled THP1 cells for another 1 hour. Following washing, attached THP1 cells were visualized and counted on an inverted fluorescent microscopy. Scale bars 1000 μm. n=6. Significance was determined by a 2-way ANOVA with Bonferroni’s post hoc test (**C** and **E**).

To define the *in vivo* therapeutic effects of XMU-MP1, we again employed a LPS-induced murine ALI model ^22^. In this regard, mice were pretreated with XMU-MP1 (4 mg/kg, IP) for 2 hours before LPS instillation. 24 hours after LPS administration, BALF and lung tissues were collected to evaluate the therapeutic effects of XMU-MP1 on ALI. As shown in Figure 6A and 6B, instillation of LPS resulted in a dramatic increase in BALF neutrophil and protein levels in vehicle treated mice, which was markedly inhibited in XMU-MP1 treated mice. Likewise, XMU-MP1 treatment substantially attenuated LPS-induced expression of chemokines and cytokines in both BALF and lung tissues, as determined by qPCR and ELISA (Figure 6C and D). LPS-induced myeloperoxidase (MPO) activity, which is a measure of neutrophil activity and an indirect indicator of lung injury, was also inhibited by XMU-MP1 treatment. Pulmonary vascular leakage, as determined by Evans blue dye extravasation, was similarly decreased in XMU-MP1 treated mice. Furthermore, our pilot studies showed that administration of XMU-MP1 after LPS instillation significantly decreased total protein levels, inflammatory cells, and MCP1 levels in BALF, while the levels of IL-6 and TNF-α only trended towards reduced levels (Figure S3). Future studies will be focused on optimizing treatment schedule and testing therapeutic efficacy in other lung injury models, such as bacterial pneumonia models. Taken together, these data suggest that pharmacological inhibition of Mst1/2 can effectively prevent EC activation and LPS-induced ALI in mice and points to a role for Mst1 inhibitor therapies in the management of inflammatory vascular diseases of the lung.

**Figure 6.**
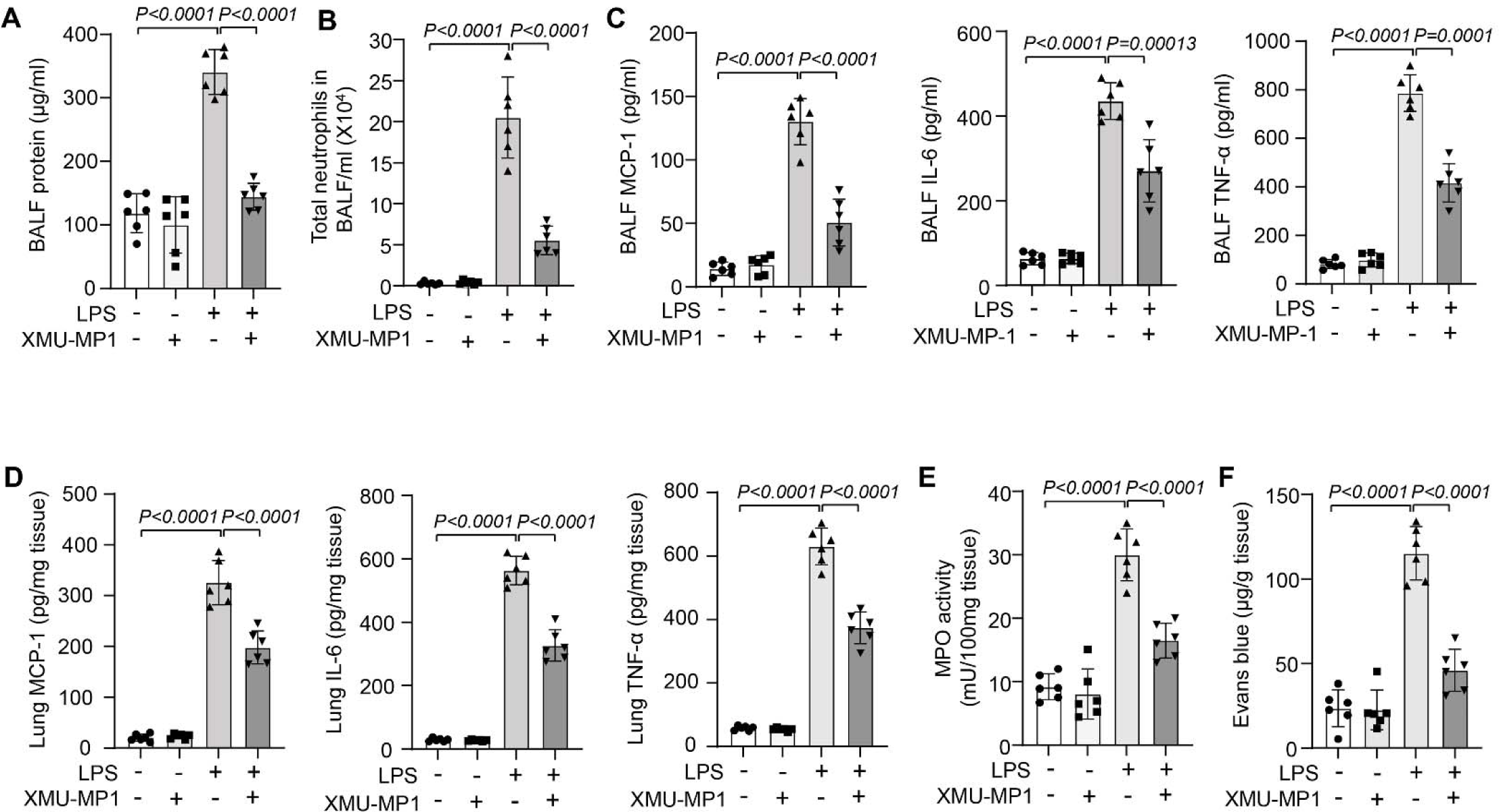
Pharmacological pretreatment to inhibit Mst1 attenuates LPS-induced ALI in mice. WT C57BL6 mice were pretreated with either vehicle or XMU-MP1 (4 mg/kg, IP) for 2 hours and then subjected to LPS instillation. 24 hours after LPS instillation, total protein (**A**), neutrophils (**B**), and the levels of MCP-1, IL-6, and TNF-α (**C**) in bronchoalveolar lavage fluid (BALF) at baseline and after LPS stimulation were determined. **D**, The levels of MCP-1, IL-6, and TNF-α in lung samples at 24 hours after LPS instillation were determined by ELISA. The activity of MPO (**E**) and Evans blue dye extravasation (**F**) in lung of vehicle and XMU-MP-1 treated mice were determined. n = 6/group. Significance was determined by a 2-way ANOVA followed by a Sidak multiple comparison test (**A-E**).

## Discussion

The Mst1/2 are core members of the Hippo pathway ^36^ and have been implicated in the pathobiology of various diseases ^37^. For instance, recent studies have demonstrated that Mst kinase is critically involved in the development of several lung diseases, such as pulmonary artery hypertension (PAH) ^38^, lung fibrosis ^39 40^, pneumonia ^41^, and lung regeneration ^20^. However, the importance of Mst kinase in controlling acute lung injury has not been investigated. In the present study, we show that Mst1 is activated in lung ECs in response to TNF-α stimulation and that activation contributes to heightened endothelial responses by increasing NF-κB signaling. Moreover, we demonstrate that genetic and pharmacological approaches to inhibiting Mst1 activity render mice more resistant to LPS-induced lung injury, supporting the notion of Mst1 directed therapies for the treatment of human disease.

The lung endothelium serves as a formidable barrier against the influx of cells and the leakage of proteins and other circulating materials in the lung ^2^. It has been increasingly recognized that the pulmonary endothelium plays a pivotal role in orchestrating inflammatory responses to pulmonary insults ^2^. In response to a range of pathological insults, such as hypoxia, cytokines, chemokines, and bacterial endotoxins, the lung endothelium produces a variety of cytokines and leucocyte adhesion molecules including intracellular adhesion molecule-1 (ICAM-1), vascular cell adhesion molecule-1 (VCAM-1) and E-selectin, which helps to recruit inflammatory cells from the circulation and support their transendothelial migration into the lung parenchyma ^7^. Thus, therapeutic strategies to suppress endothelial inflammation are anticipated to ameliorate ALI and other inflammatory vascular conditions of the lung. At the molecular level, activation of NF-κB is fundamentally important in driving endothelial inflammation and the upregulation in expression of endothelial cell adhesion molecules. Upstream regulators of NF-κB in ECs have been poorly identified and require further investigation. Mst1 is an important component of the hippo/yap pathway that has been implicated in organ size control, tissue regeneration, and self-renewal ^36^. Mst1 is proteolytically cleaved by caspase to 34-36-kDa N-terminal catalytic fragment, which increases Mst1 kinase activity and nuclear translocation ^42 43^. In addition to caspase cleavage, Mst1 phosphorylation at Thr183 has been proposed to contribute to kinase activation ^32^. In accordance with these findings, we found that short- and long-term exposures to TNF-α led to a significant increase in Mst1 phosphorylation at Thr183 and enhanced Mst1 cleavage, which constitutes a potent feedback loop in Mst1 activation, further highlighting the critical role that sustained Mst1 activation has on lung EC inflammation.

Recently, several studies have implicated the regulatory roles of Mst1 in NF-κB activation ^16 44 45 46 47^. Depending on the cell type, Mst1 has been shown to promote or inhibit NF-κB activation. For instance, Mst1 has been shown to promote NF-κB activation in brain tissues, cardiomyocytes, and skin lesions of psoriasis ^16 46 48^. On the other hand, Mst1 activation inhibits NF-κB activation in fibroblasts and cancer cells ^45 49^. The underlying reasons to explain these opposing effects have not been explained. In this study, we found that Mst1 activation promotes NF-κB activation in lung ECs and that overexpression of DN-Mst1 suppressed NF-κB activity, supporting the notion that endogenous Mst1 regulates the transcriptional activity of NF-κB in lung ECs. We and others have shown that Mst1 is an upstream kinase of the JNK and p38 MAPK pathways ^50 51^, which have been shown to induce NF-κB activation in a wide range of cells ^52^. Consistent with these observations, we found that Mst1 induces the activation of the JNK and p38 MAPK pathways in lung ECs and that pharmacological inhibition of JNK and p38 MAPK markedly attenuated Mst1-induced NF-κB activation in ECs, further suggesting the role of the JNK and p38 MAPK in Mst1-induced NF-κB activation in lung ECs.

XMU-MP-1 has been identified as a potent and specific inhibitor of MST1/2 kinases and it has been shown to inhibit MST1/2 activities in various *in vitro* and i*n vivo* studies ^35 53 54^. In this study, we demonstrated that pharmacological inhibition of Mst1 by XMU-MP-1 dramatically suppressed lung inflammation and vascular leakage to LPS in mice. Further, our *in vitro* studies showed that XMU-MP-1 dramatically attenuated TNF-α-induced cytokines, chemokine, and VCAM-1 expression. Interestingly, treatment did not have a significant effect on ICAM-1 mRNA levels, but significantly inhibit ICAM-1 protein expression in lung ECs, suggesting that XMU-MP-1 may regulate ICAM-1 expression at post-translational levels, which needs further investigation.

The decision to administer XMU-MP-1 prior to the induction of ALI points to a role for Mst1 inhibitor treatments in the prevention of ALI. We speculate this could be important for patients at high risk for ALI and ARDS. Further, our recent pilot studies demonstrated that administration of XMU-MP-1 after LPS exposure also inhibited lung permeability and the expression of MCP-1 in BALF, indicating a potential therapeutic capacity of this inhibitor for established disease. Future studies will be needed to optimize the therapeutic regimen and test the efficacy of Mst1 inhibitors in other lung injury models.

In summary, our results provide both in vitro and in vivo evidence to demonstrate that Mst1 is a critical regulator of lung endothelial activation and inflammation through activation of the NF-κB pathway. Targeted inhibition of Mst1 may represent a novel therapeutic strategy for the prevention and treatment of inflammatory lung diseases, such as ALI and ARDS.

## Author contributions

ZG, NT, CL, CZ, LH, DC, and MMZ performed experiments, ZG, NT, CL, CZ, RS, and JS designed experiments, analyzed and interpreted data, ZG, RS, and JS wrote the paper, all authors edited the paper.

## Sources of Funding

This work was funded by National Heart, Lung, and Blood Institute grant R01HL159168 and R01HL152703 (R.S. and J.S.), and American Heart Association Established Investigator Award 16EIA27710023 (J.S.).

## Disclosures

The authors have declared that no conflict of interest exists.

## Notes

### Competing Interest Statement

The authors have declared no competing interest.

